# Par3/Baz levels control epithelial folding at actomyosin-enriched compartmental boundaries

**DOI:** 10.1101/125500

**Authors:** Jose M. Urbano, Huw W. Naylor, Elena Scarpa, Leila Muresan, Bénédicte Sanson

## Abstract

Epithelial folding is crucial to shape embryos and tissues during development. Here we investigate the coupling between epithelial folding and actomyosin-rich boundaries. The mechanistic relationship between the two is unclear, since actomyosin-rich boundaries can be either associated with folds or not, while epithelial folding has been found to be either dependent or independent of actomyosin contractility. Here we investigate the shallow folds that form at compartmental parasegment boundaries (PSBs) in the early *Drosophila* embryo. First, we demonstrate that formation of these folds is dependent on the contractility of supracellular actomyosin cables. When the Myosin II phosphatase Flawing is depleted at the PSBs, actomyosin contractility increases, resulting in deeper folds. Conversely, in *wingless* mutants, actomyosin enrichment and increased contractility at PSBs are lost and this correlates with an absence of folding. Furthermore, when we make ectopic PSBs by expressing Wingless ubiquitously, the ectopic boundaries become enriched in actomyosin and epithelial folds form. Ectopic PSB folds, however, are much deeper than endogenous ones, indicating that epithelial folding is normally under inhibitory control. We present evidence that depletion of Bazooka/Par-3 levels at PSB cell-cell contacts, which is under Wingless signaling control, is responsible for this inhibition. Bazooka is found depleted at endogenous but not ectopic PSBs. In embryos overexpressing Bazooka, endogenous PSB folds form earlier and are much deeper. To ask how local signaling at the boundaries control Bazooka levels at cell-cell contacts, we examined embryos that ectopically expressed Wingless in an *hedgehog* mutant background. In these embryos, inhibition of folding is rescued, with ectopic PSBs now forming shallow folds as endogenous PSBs. Bazooka is depleted at these ectopic PSBs in absence of Hedgehog, suggesting an opposite effect of Wingless and Hedgehog signaling on Bazooka levels at PSB cell-cell contacts. This uncovers a new role of Bazooka in controlling fold formation at actomyosin-rich compartmental boundaries.

## Introduction

Epithelial sheet bending is ubiquitous in animal development and essential to elaborate the complex anatomy of the body. It is observed throughout development, from the invaginations and involutions of gastrulation to the folding of organ tissues (Keller and Shook, 2011; Bazin-Lopez et al., 2015). The mechanisms identified so far that promote epithelial sheet bending are diverse (Pearl et al., 2017). One of the best studied mechanism is apical constriction mediated by actomyosin activation at the apical end of epithelial cells (Martin and Goldstein, 2014). Yet not all mechanisms uncovered for epithelial folding require actomyosin activity: for dorsal folds formation in *Drosophila* embryos, a basal shift of the adherens junctions is required instead (Wang et al., 2012).

Here we investigate the relationship between epithelial folding and planar actomyosin cables that are often found in developing epithelia (Monier et al., 2011; Roper, 2013; Fagotto, 2015). For example, actomyosin enrichments at the level of adherens junctions line up compartmental boundaries in *Drosophila* epithelial tissues, where they are required for lineage restriction (Landsberg et al., 2009; Monier et al., 2010). In some instances, such as in segmented tissues, actomyosin-rich cables are associated with folds (Mulinari et al., 2008; Calzolari et al., 2014). In other cases, such as for the antero-posterior (AP) compartmental boundary in *Drosophila* wing discs, actomyosin-rich boundaries are anatomically “silent”, with no folding observed (Landsberg et al., 2009). Intriguingly, however, some mutant backgrounds can generate folds at the AP boundary in wing discs, indicating that fold formation is normally suppressed at this compartmental boundary (Shen et al., 2008; Liu et al., 2016).

In *Drosophila* embryos, the AP compartmental boundaries (called parasegmental boundaries, PSBs) first enrich actomyosin at gastrulation, during germ-band extension (Tetley et al., 2016). Wingless signaling on one side of the compartmental boundary maintains these enrichments at germ-band extended stage, where they act as mechanical barriers to keep dividing cells in their compartment of origin (Monier et al., 2010; Monier et al., 2011). Although actomyosin enrichments are present, no epithelial folding is associated with PSBs during gastrulation and it is only towards the end of germ-band extended stage (stage 10-11) that shallow indentations form, the parasegmental grooves (Larsen et al., 2008). Here, we first establish that actomyosin contactility is indeed required for fold formation at PSBs. Next, we provide evidence that depletion of Bazooka, the homologue of vertebrate Par-3, is required to moderate fold formation at PSBs under the control of Wingless signaling. Together with a previous study at AP boundaries in the wing disc (Liu et al., 2016), this indicates that specific mechanisms exist to suppress fold formation at actomyosin-rich boundaries and our work uncovers a new role for Bazooka in this process.

## Results and discussion

### Planar polarities and increased actomyosin contractility at the PSBs require Wingless signaling

To understand what could control epithelial folding at the PSBs, we asked if the proteins known to be planar polarized at cell-cell junctions during axis extension are polarized at PSBs at germ-band extended stages. For example, during germ-band extension, F-actin, Myosin II and Rho-kinase (Rok) are enriched at dorso-ventral (DV)-oriented junctions, whereas Bazooka (Baz, Par-3 homologue) and E-cadherin are depleted (Bertet et al., 2004; Zallen and Wieschaus, 2004; Blankenship et al., 2006; Simoes Sde et al., 2010; Levayer et al., 2011). We had shown previously that F-actin and two reporters for Myosin II, MHC-GFP and MRLC-GFP, are enriched at PSBs at stage 10 (Monier et al., 2010). By quantifying the enrichment along the PSBs relative to control columns of DV-oriented junctions, we now find that Myosin II is not only enriched but also activated: the mono-phosphorylated form of MRLC (recognized by the Sqh1P antibody, (Zhang and Ward, 2011) accumulates at the PSB (Fig. 1 A-B’, E). Using the same quantification method, we find that Rok (labelled using the reporter rok-GFP) is enriched at the parasegment boundary while Baz is depleted (Fig. 1 E and Fig. S1 B-C’), as seen in DV-oriented junctions at gastrulation. Therefore the key planar polarities of germ-band extension are maintained at germ-band extended stage specifically at the PSBs (the remaining of the epithelium does not show any detectable planar polarities). We find however one notable difference in our survey of polarities: whereas E-Cadherin levels are depleted at DV junctions in germ-band extension (Blankenship et al., 2006; Levayer et al., 2011), this is not the case at PSBs, where levels are the same as in other junctions (Fig. 1E and Fig. S1 A-A”). This is interesting as this might be linked to the distinct behaviours of cells during axis extension versus boundary formation: when cells intercalate during GBE, contacts need to be broken and the depletion of E-Cadherin might facilitate this. In contrast, normal adhesion might be required for boundary formation at later stages.

**Figure 1:**
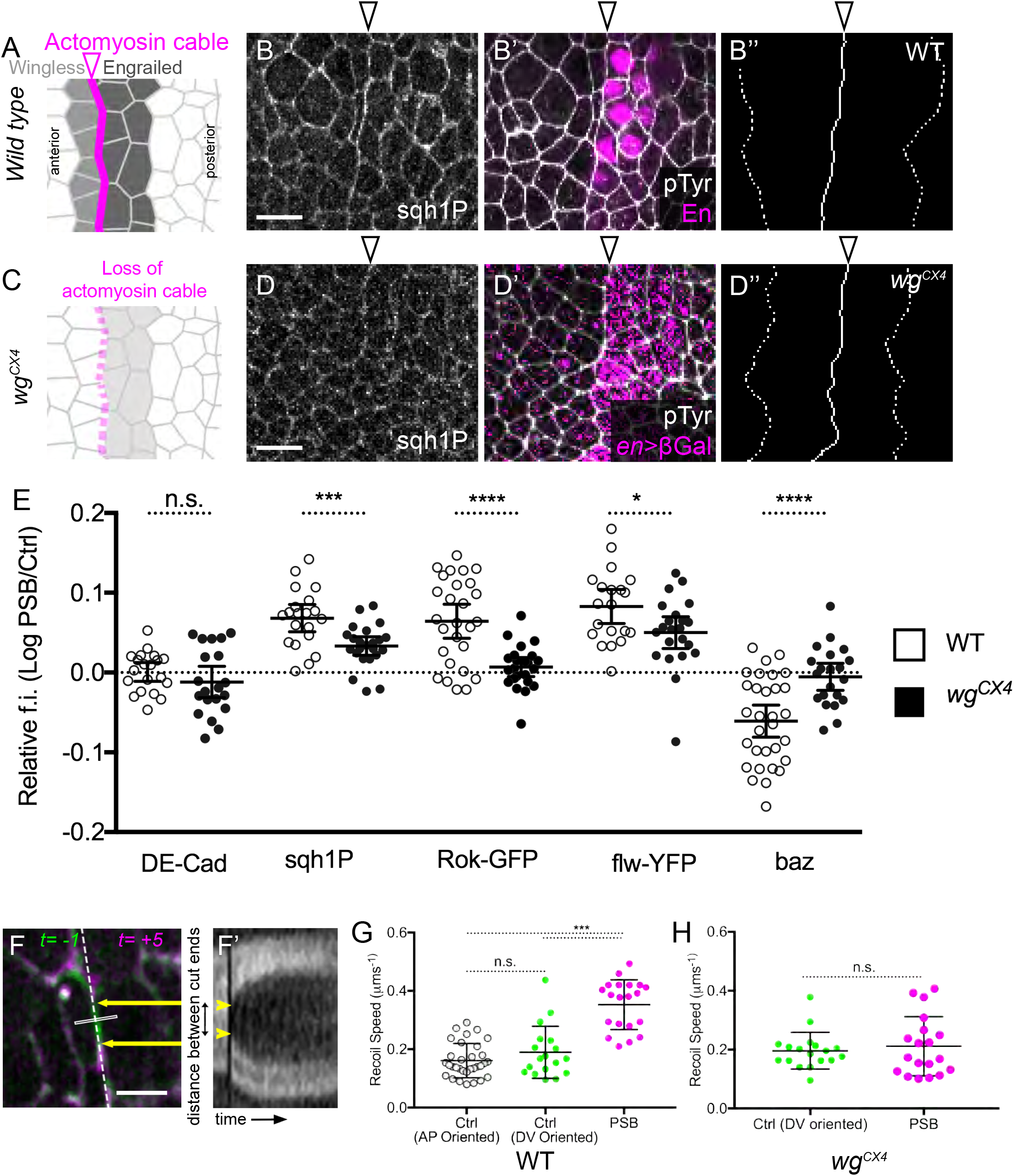
Planar polarities and actomyosin contractility at the PSBs in WT and *wingless* mutant embryos. (A) Diagram showing the position of the actomyosin cable at the parasegmental boundary (PSB) located at the interface between *wingless* and *engrailed*-expressing cells in wild-type embryos of stage 9/10. In *wingless* mutant embryos (C), the actomyosin cable is lost. Immunostainings against Sqh1P (B, D) and pTyr and Engrailed or ß-gal (B’, D’) showing the presence and absence of Myosin II enrichment at the PSBs, respectively, in wild type (B-B”) and *wingless* mutant early stage 10 embryos (D-D’). In *wingless* mutants, *engrailed* expression is not maintained (C), so a *en-lacZ* reporter was used to trace *engrailed* expression and thus the PSB location (Vincent and O’Farrell, 1992). (B”, D”) Corresponding tracings of the PSB (continuous line) and control cell-cell contacts (dotted line) to quantify the Sqh1P signal. (E) Quantification of the enrichment of different markers at the PSB relative to control cell interfaces, in wild type and *wingless* mutant embryos. (F-H) Laser ablations to probe junctional tension at the PSBs. (F) Overlay before (green) and after (pink) ablation of a single cell-cell junction at the PSB (white rectangle: ablation line). Scale bar = 5 mm. (F’) A kymograph of the signal along the dashed line in F was used to measure the distance between the recoiling cut ends over time (yellow arrows and arrowheads). Speed of recoil upon ablation of cell-cell junctions in wild type (G) and *wingless* mutant embryos (H), either at PSBs or control junctions oriented parallel to the antero-posterior (AP) or dorso-ventral (DV) embryonic axes (PSBs are DV-oriented). In all figures anterior is left and dorsal up. Empty arrowheads depict PSBs.

Whereas planar polarities during germ-band extension are under the control of the pair-rule genes (Bertet et al., 2004; Zallen and Wieschaus, 2004), we have shown previously that robust actomyosin polarization at PSBs at germ-band extended stage (stage 10) requires Wingless signaling (Monier et al., 2010). Confirming this, the enrichment of Sqh1P is significantly decreased at PSBs in *wingless* null mutants (Fig. 1 C-E). Moreover, the enrichment of Rok and depletion of Baz at PSBs is lost (Fig. 1 E). Therefore the planar polarities observed at the PSBs require Wingless signaling. It also suggests that Wingless signaling increases actomyosin contractility along the PSBs. We have provided previously some evidence of this by showing that the PSBs are straighter than control columns of interfaces, an indication of higher tension along the PSBs, and that this increased straightness is lost in *wingless* mutants (Monier et al., 2010). Here we probe more directly junctional tension at the PSBs, using laser ablation to measure the speed of junctional recoil as previously (Tetley et al., 2016). We find that, at stage 10, the initial velocities of recoil are about twice as fast at PSBs compared to non-boundary DV or anterior/posterior (AP)-oriented junctions (Fig. 1 F-G and Fig. S1 E, E’). This difference is lost in *wingless* mutants (Fig. 1 H and Fig. S1 F, F’). This demonstrates that Wingless signaling is required for increasing actomyosin contractility and junctional tension at the PSBs at germ-band extended stages.

### Hyperactivation of Myosin II via knockdown of the Myosin II phosphatase Flw increases epithelial folding at the PSB

Since parasegmental groove formation also requires Wingless signaling (Larsen et al., 2008), the above results suggest that epithelial folding at PSBs is a direct consequence of increased actomyosin contractility. To test this, we increased actomyosin contractility further by knocking down the Myosin II phosphatase, Flapwing (Flw) (Vereshchagina et al., 2004). Using a CPTI protein trap line with YFP inserted in the *flw* locus (Lowe et al., 2014; Lye et al., 2014), we show that Flw-YFP is enriched at the PSBs and this enrichment is significantly reduced in *wingless* null mutants (Fig. 1E and Fig. S1 D-D”). This suggests that Myosin II activity is kept in check at the PSB by Flw. To disrupt this negative regulation, we used the nanobody system to degrade Y/GFP-tagged proteins (Caussinus et al., 2012). Since Flw-YFP is homozygous viable, all molecules of Flw are susceptible to be degraded. When nanobodies against YFP (*UAS-deGradFP*) are expressed under the control of *paired-Gal4* (*prd-Gal4*), Flw-YFP is efficiently depleted in every other parasegment (Fig. 2A). Whereas Flw-YFP is normally mainly cortical, it is lost from the membranes in the *prd-Gal4* domains and accumulates as bright dots in the cytoplasm (Fig. 2B). In those domains, Sqh1P levels are elevated (Fig. 2B-B”), indicating that Flw depletion by nanobodies results in Myosin II hyperphosphorylation as in *flw* mutants (Vereshchagina et al., 2004; Sun et al., 2011). Not only is this hyperactivation detectable throughout the *prd-Gal4* domains, there is also a higher enrichment of Sqh1P specifically at the PSBs (Fig. 2F). We note that Baz levels remains depleted at those PSBs (with some increase in the level of depletion) and E-cadherin is unchanged (Fig. 2F). The Myosin II hyperactivation at the PSBs results in a specific morphogenetic phenotype: the normally shallow parasegmental grooves (Fig. 2C) become very deep in every *prd-Gal4* domains (Fig. 2D, E and Fig. 2G-H”). This effect is specific to the PSBs: no other epithelial folds appear in the domains depleted for Flw. This suggests that epithelial folding is a tightly controlled mechanism that happens at competent cell interfaces such as the PSBs. Together, these results indicate that increasing actomyosin contractility at the PSBs is sufficient to enhance epithelial folding.

**Figure 2:**
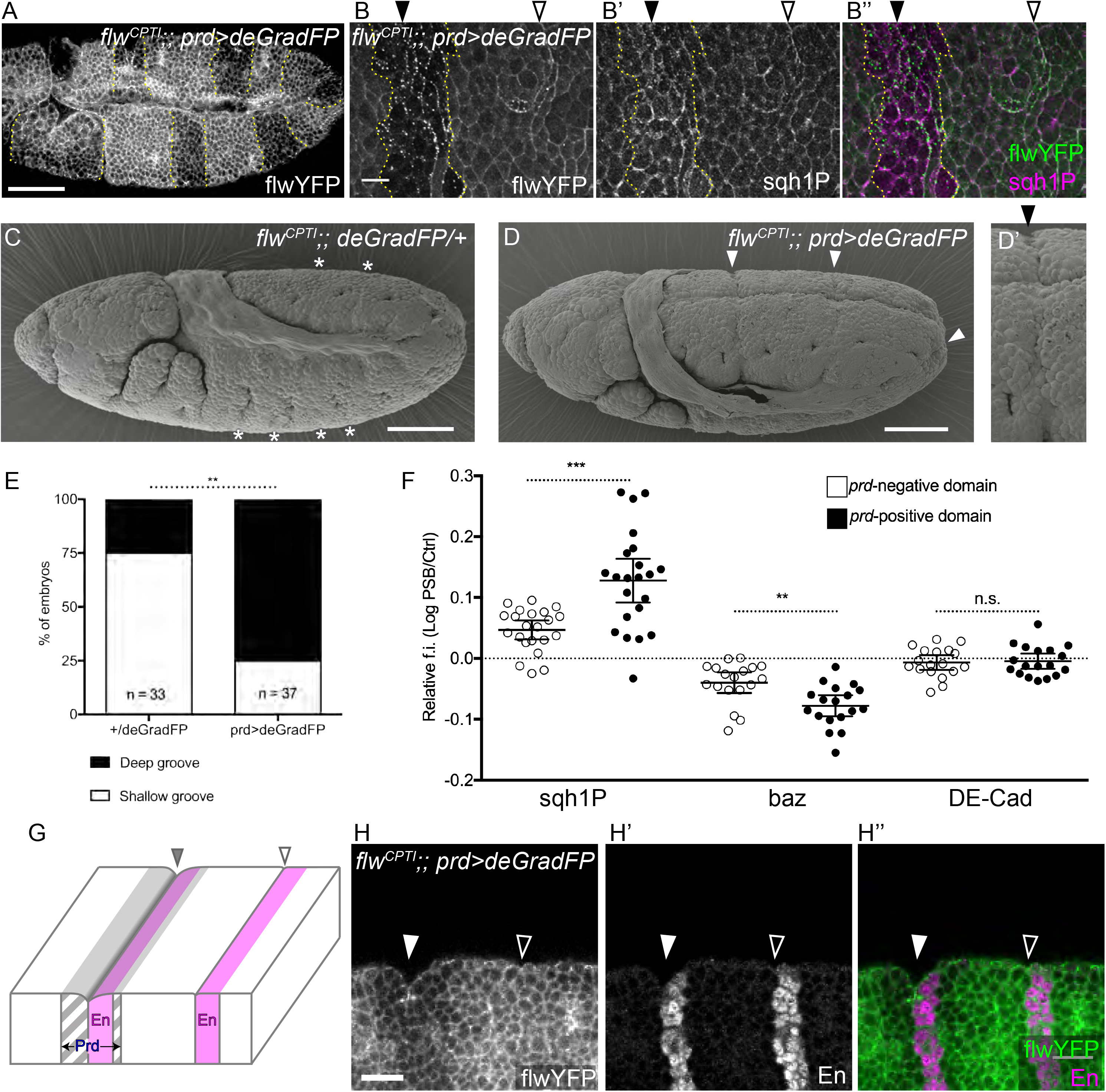
PSB grooves are deepened by depletion of Flw. (A) Immunostaining against GFP to reveal the degradation of Flw-YFP in a stage 10 embryo expressing the DeGrad nanobodies against GFP (deGradFP) in the *prd-Gal4* domain (highlighted with yellow dotted lines). (B-B”) Embryos of the same genotype at higher magnification, immunostained against GFP (B), and Sqh1P (B), with merge in (B”). (C, D) Scanning electron microscopy (SEM) of a stage late 10 control (C) and deGradFP (D) embryo. The parasegmental grooves are labelled with stars: these are very shallow indentations spanning the whole DV width of the germ-band in control embryos (C). By contrast, in embryos where Flw is depleted in *prd-Gal4* domains, the parasegmental grooves deepen (arrowheads) (close up in D’). (E) Blind quantification of the number of embryos with either shallow only or deep groves in the sibling embryos shown in C and D. (F) Quantification of the enrichment of Sqh1P, Baz and E-cad at the PSB in the deGradGFP expressing and non-expressing domains (Prd-Gal4 positive or negative). (G) Schema showing the position of the different type of parasegmental grooves in relation to Engrailed and the *prd-Gal4* expression domains. (H-H”) Immunostaining against GFP (H) and Engrailed (H’) (H”, merge) to show the deep groove at the PSB in the Flw depleted domain (solid arrowhead). The wild type PSB position is indicated (empty arrowhead).

### Increased epithelial folding at ectopic PSBs in embryos overexpressing *wingless*

To further investigate the link between actomyosin contractility and epithelial folding we generated ectopic parasegment boundaries by expressing *wingless* in the whole epithelium (*arm-Gal4/UAS-wg*, hereafter *arm>wg*) (Sanson et al., 1999; Larsen et al., 2008). In these embryos, Engrailed expression expands posteriorly to fill its competence domain and an ectopic boundary forms at the posterior margin of the Engrailed expressing cells (Fig. 3A). This unique site replicates the signaling environment of the endogenous parasegmental boundary, with Hedgehog being able to signal from the Engrailed competence domain into the now adjacent Wingless competence domain. As for endogenous PSBs, we find that Sqh1P is enriched at the cell-cell contacts of the ectopic PSBs relative to control interfaces (Fig. 3B-B” and C). The positive and negative regulators of Myosin II, Rok and Flw, are also enriched at the ectopic PSBs (Fig. 3C). Furthermore, laser ablations of cell-cell contacts at ectopic PSBs show that junctional tension is elevated there as for endogenous PSBs (Fig. 3D and Fig. S2 B,B’). We conclude that ectopic PSBs exhibit increased actomyosin contractility and junctional tension as endogenous PSBs.

**Figure 3:**
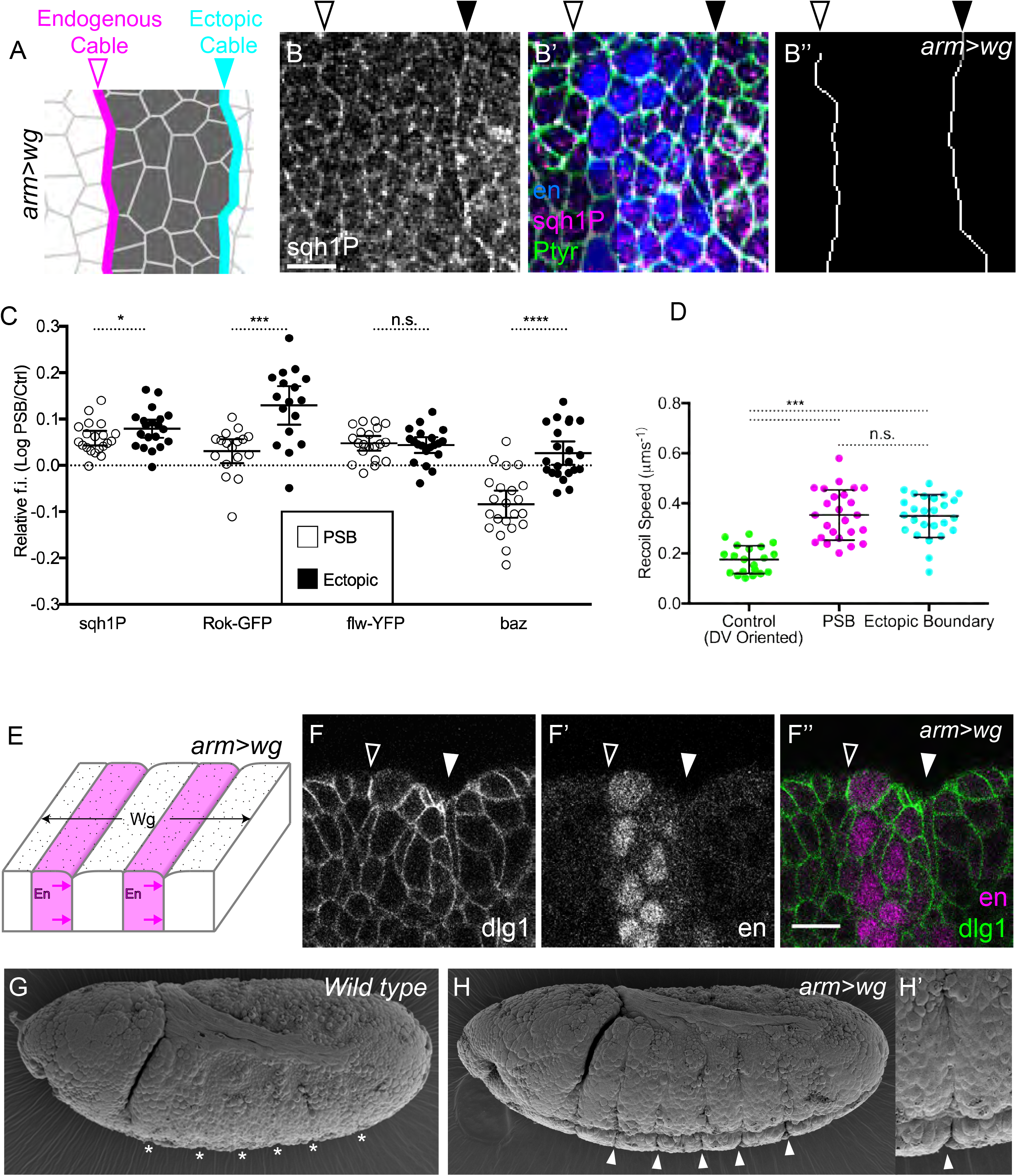
Increased epithelial folding at ectopic actomyosin boundaries in *wingless-*overexpressing embryos. (A) Diagram showing the position of the endogenous (pink) and ectopic (blue) parasegmental boundaries, in *arm-Gal4/UAS-wg (*shorten to *arm>wg)* embryos. The ectopic PSB forms at the posterior interface of the enlarged Engrailed domain (dark cells), in each parasegmental repeats. (B-B”) Immunostaining of *arm>wg* early stage 10 embryos against Sqh1P (B), Engrailed and pTyr (merged in B’). (B”) Traces of the endogenous (empty arrowheads) and ectopic (solid arrowheads) PSBs. (C) Quantification of the enrichment of planar polarized markers at the endogenous (empty circles) and ectopic (solid circles) PSBs, relative to control DV-oriented interfaces. (D) Initial recoil speeds for laser ablation of endogenous and ectopic PSB cell junctions, and control DV-oriented cell interfaces. (E) Diagram of the position of the shallow and deep folds at endogenous and ectopic PSBs in *arm>wg* embryos. (F-F”) View of the curvature of an *arm>wg* embryo stained for Dlg (F) and Engrailed (F’) (merge in F”) to show the difference in depth between endogenous (empty arrowheads) and ectopic (solid arrowheads) PSBs. Scanning electron microscopy of stage 9 wild type and *arm>wg* embryos. Endogenous PSBs are barely indenting the surface of the embryo (G) while ectopic PSBs form deep grooves (H) (close-up in H’).

Ectopic boundaries are also associated with an epithelial fold, providing additional evidence that actomyosin contractility at PSBs promote epithelial folding. There is however a key difference: folds at ectopic PSBs are deep and form earlier in development when compared to endogenous parasegmental grooves (Fig. 3 E-H’). The folding seems even more pronounced to what is observed when actomyosin contractility is elevated in the Flw knock-down (Compare Fig. 2 D and Fig. 3 H, H’). However, in the case of the ectopic PSBs, we cannot find evidence of a further increase in actomyosin contractility compared to the endogenous PSBs, which could explain the deep folds: enrichments of Sqh1P at ectopic and endogenous PSBs are similar (Fig. 3C) and the recoil speeds upon laser ablation are undistinguishable (Fig. 3D). We examined ectopic PSBs in two other genotypes, *rho-Gal4/UAS-wg (rho>wg)* and null mutants for the gene *naked* (*nkd*). *Rho-Gal4* is expressed in a ventral stripe a few cell diameters wide on either side of the ventral midline (Ip et al., 1992). When *wingless* is ectopically expressed using this driver, the Engrailed domain is enlarged in the corresponding ventral region and a ventral, stubby, ectopic PSB forms in each parasegment, which is enriched in actomyosin (Fig. S2 C, E-E”). Despite its short length, a deep fold forms at this ectopic PSB (Fig. S2 D, D’). The fold does not extend beyond the site of the ventral ectopic PSB (visualized by Engrailed staining and Sqh1P enrichment, Fig. S2 E-E”), suggesting that the increased folding is cell-autonomous. In a *nkd* embryo, Wingless signaling is altered and signals more weakly but further, resulting in an enlarged Engrailed domain and an ectopic PSB (Martinez Arias et al., 1988; Zeng et al., 2000). These ectopic PSBs enrich actomyosin and produce a deep fold (Fig. S2 F-H”). Thus, ectopic PSBs produced by different genetic manipulations are consistently associated with increased epithelial folding.

We looked for factors that could explain the difference in the degree of epithelial folding between endogenous and ectopic PSBs. We note that Rok is more enriched at ectopic PSBs than endogenous ones (Fig. 3C). However, since Sqh1P enrichment (also Sqh-GFP enrichment, Fig. S2 B) and junctional tension are similar at endogenous versus ectopic PSBs (Fig. 3 C, D), the increase in Rok is unlikely to have an impact on actomyosin contractility at the ectopic PSBs (but it could affect folding through other pathways). The other difference we find is that Baz is not depleted at ectopic boundaries (Fig. 3 C and Fig. S2 A-A”). The fact that Baz/Par-3 has been implicated in the initiation of dorsal folds in the early embryo (Wang et al., 2012) prompted us to analyze a putative role of Baz/Par-3 in controlling epithelial fold depth at parasegmental boundaries.

### Bazooka/Par-3 increases the depth of epithelial folding at PSBs

To test if Baz could have an impact on epithelial folding at PSBs, we overexpressed UAS-Baz-GFP in the embryo using maternal Gal4 drivers (*MTD>bazGFP*, quantified in Fig. 4D). We find that Baz overexpression causes the formation of deep folds specifically at the PSBs and nowhere else (Fig. 4A-B”). These folds are much deeper and also form earlier than wild-type parasegmental grooves. As for ectopic PSBs in *arm>wg* embryos, this effect cannot be explained by an increase in actomyosin contractility. First, laser ablations of PSB versus control DV-oriented junctions give similar initial recoil velocities (Fig. 4C, compare with Fig. 1G or Fig. 3D), with a similar ratio of ~2 between PSBs and control junctions (Fig. 4C and Fig. S3 B, B’). Second, the absolute quantities of Sqh1P are equivalent between Baz overexpressing embryos and wild type, for both PSB and DV-oriented control junctions (Fig. 4E). Third, Sqh1P is similarly enriched at PSBs in both genotypes (Fig. S3 A). So in term of actomyosin contractility and junctional tension, the PSBs in Baz overexpressing embryos are undistinguishable from those in wild type embryos. Interestingly, Baz is still found depleted at PSBs compared to control junctions in Baz overexpressing embryos (Fig. S3 A), suggesting that the signals controlling its depletion at boundaries are functioning normally. As expected, however, the absolute levels of Baz are much higher in Baz overexpressing embryos (Fig. 4D), indicating that it is the overall increase in Baz that promotes deep epithelial folding at actomyosin-enriched boundaries.

**Figure 4:**
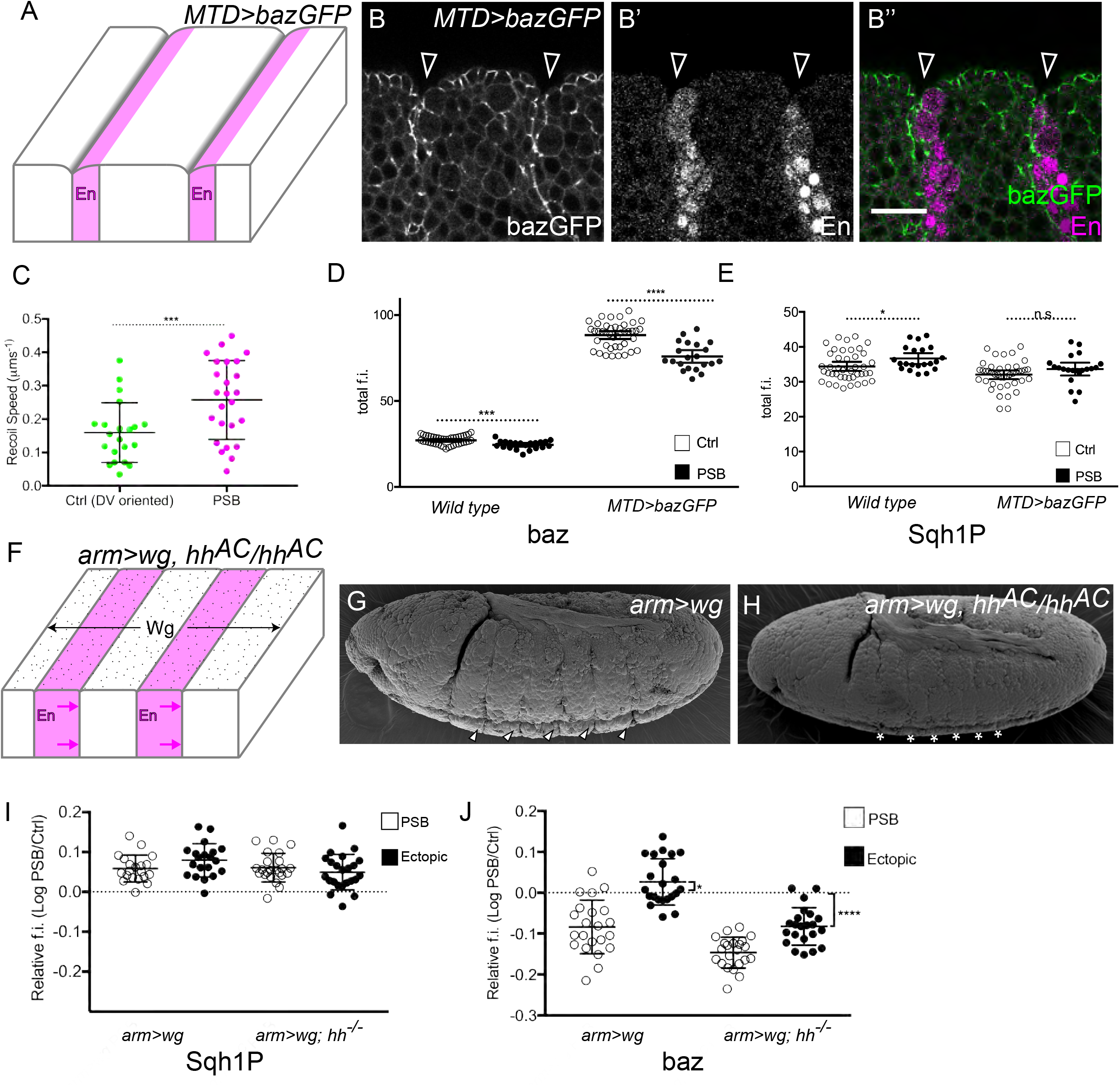
Impact of Baz on the depth of epithelial folding at actomyosin boundaries. (A) Diagram showing the position of deep folds at endogenous PSBs in embryos ectopically expressing Baz (*MTD>bazGFP* embryos). (B) Immunostaining of these embryos at early stage 10 against GFP (B) and Engrailed (B’), merged in (B”), showing that deep grooves form at PSBs. (C) Speed of recoil upon ablation of cell-cell junctions in *MTD>bazGFP* embryos, either at PSBs or control junctions. (D) Quantification of absolute quantities of baz (D) and Sqh1P (E) at the PSBs and control DV-oriented cell interfaces in wild type (empty circles) and *MTD>bazGFP* (solid circles) embryos. (F) Diagram showing the lack of deep grooves at endogenous and ectopic PSBs, in *arm>wg* embryos with a null mutant background for *hh*. SEM images showing that the deep grooves at ectopic PSBs (arrowheads) seen in stage 9 *arm>wg* embryos (G) become shallow (stars) in a *hh* null mutant background (H). Quantification of the enrichment of Sqh1P (I) and Baz (J) at the endogenous and ectopic PSBs relative to control DV-oriented cell interfaces, in stage 10 *arm>wg* and *arm>wg; hh^AC^/hh^AC^* mutant embryos.

To test this further, we searched for experimental conditions that could rescue deep epithelial folding. We showed that Wingless signaling is required for Baz depletion at the endogenous PSBs (Fig. 1), but Baz is not depleted at ectopic boundaries in *arm>wg* embryos (Fig. 3). So a possibility was that a signal was inhibiting Wingless-dependent depletion of Baz at ectopic boundaries. A likely signal is Hedgehog (Hh): it has been found to antagonize the regulation of specific genes by Wingless in the region posterior to the Engrailed domain (Sanson et al 1999) and to increase the lysosomal degradation of Wingless in this region (Dubois et al., 2001). This antagonistic relationship is important for segment polarity because it establishes an asymmetry in Wingless activity on either side of the parasegmental boundary (active anterior to it/inhibited posterior to it) (Hatini and DiNardo, 2001; Sanson, 2001). To test if Hedgehog signaling could have the opposite effect of Wingless signaling on Baz levels, we quantified Baz at ectopic boundaries in *arm>wg* embryos in a null mutant background for *hedgehog* (*arm>wg [hh^-/-^])*. In these embryos, fold depth is reduced and similar to those at endogenous boundaries (Fig. 4F-H). We find that Baz depletion at ectopic boundaries in *arm>wg [hh^-/-^]* is now undistinguishable from the depletion of Baz at endogenous PSB in *arm>wg* (Fig. 4J) or wild type embryos (Fig. 1E). This is consistent with the known symmetric activity of Wingless in equivalent embryos (Sanson et al., 1999; Dubois et al., 2001) and provides additional evidence that Wingless depletes Baz levels at the endogenous and ectopic PSBs. Interestingly, the depletion of Baz at endogenous boundaries in *arm>wg [hh^-/-^]* embryos is enhanced (Fig. 4J), further supporting the idea that Wingless and Hedgehog signaling have opposite effects on Baz levels at the PSBs. Together, these experiments indicate that Baz levels control the extent of epithelial folding at actomyosin-rich boundaries.

### Bazooka/Par-3 activity in epithelial folding at PSBs is independent of apical constriction or adherens junctions lowering

Known mechanisms for groove formation include apical constriction (Martin and Goldstein, 2014) and adherens junctions (AJs) lowering (Wang et al., 2012). We asked if epithelial folding at PSBs required any of these two possible mechanisms. To test this, we imaged the whole cell volume labeling membranes with actin phalloidin in WT (Fig. 5 A, A’) and *arm>wg* (Fig. 5 B, B’), in stage 9/10 embryos. We then segmented the 3D volumes of cells abutting or not the PSBs (Fig. 5 C, D). We developed a method to measure the position of the AJs relative to the apical top of the cells (Fig. 5 E, F). In both WT and *arm>wg* embryos, the AJs are a fraction lower at PSB interfaces compared to non-PSB interfaces (Figure 5 E, F). However, this lowering is not comparable with the extent of AJs basal shift observed during dorsal folding in early embryos, where the junctions of dorsal fold cells lower by up to 10 µm, while neighboring cells shift their AJs by 3µm (Wang et al., 2012). We conclude that although both involve Baz, epithelial folding at PSBs is likely to occur with a different mechanism than dorsal folding in gastrulating embryos.

**Figure 5:**
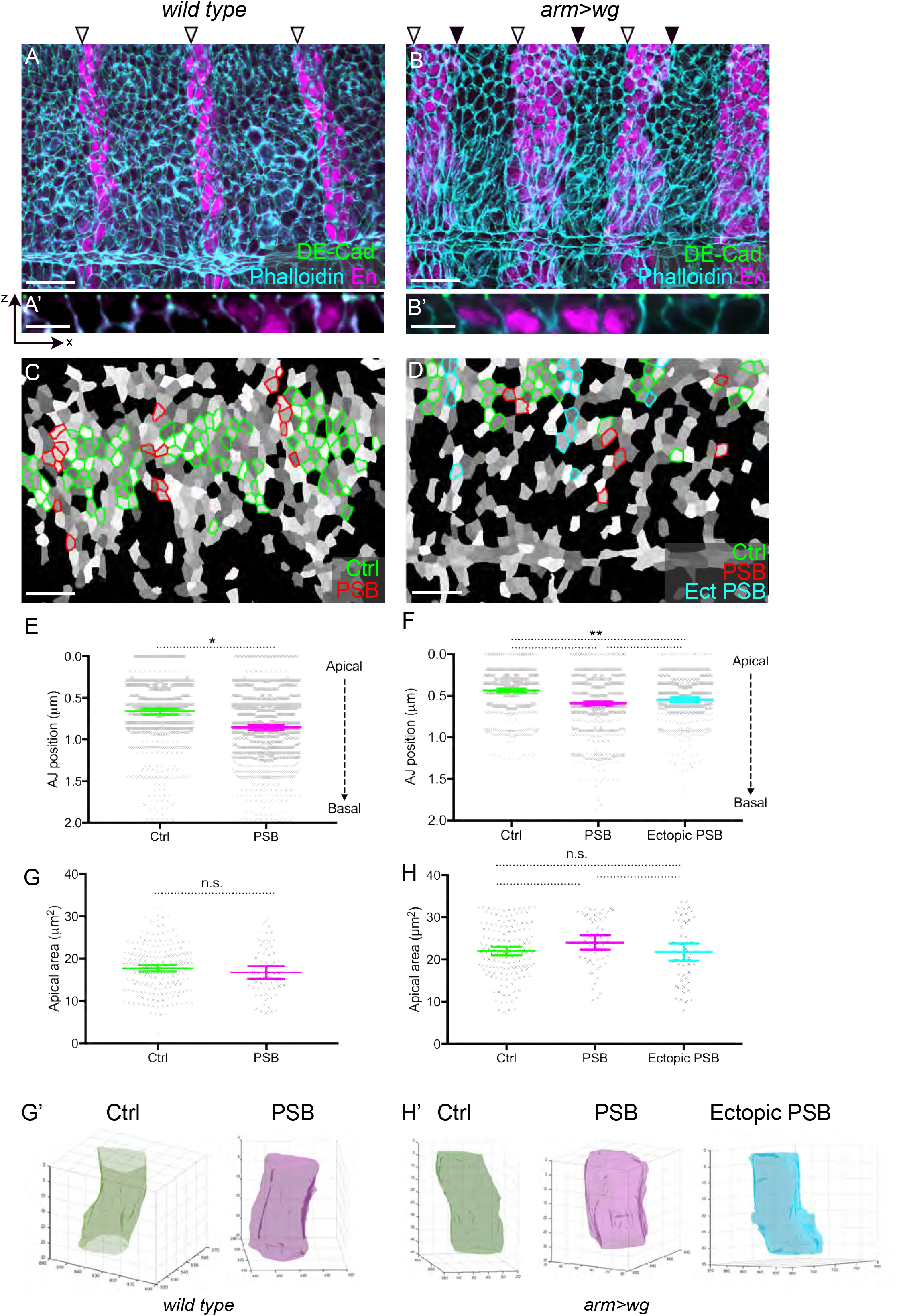
Measuring adherens junctions lowering and apical constriction in PSBs folding. (A, B) Confocal stack projections of (early stage 10) wild type and (late stage 9) *arm>wg* embryos immunostained with E-Cadherin (green), Phalloidin (cyan) and Engrailed (magenta). The positions of endogenous (empty arrowheads) and ectopic (solid arrowheads) PSBs are indicated. (A', B') X-Z optical sections of stacks shown in A and B. (C-D) Cell segmentation of image stacks shown in A and B. Cells depicted in green, red and blue represent control cells and endogenous and ectopic PSB-abutting cells, respectively. (E-F) Quantification of the distance between the position of adherens junctions (AJs) and the top of the cell for control, endogenous PSB and ectopic PSB cell-cell junctions, in wild type (E) and *arm>wg* (F) embryos. In wild type embryos (E), the mean AJ positions are -0.66 mm for control and -0.85 mm for PSBs. In *arm>wg* embryos (F), the mean AJ positions are - 0.43 mm for control, -0.59 mm for endogenous PSBs and -0.55 mm for ectopic PSBs. (G-H) Quantification of the apical area of control cells and of cells contacting either endogenous and ectopic PSBs, in wild type (G) and *arm>wg* (H) embryos. (G’,H’) 3D reconstruction of representatives of corresponding cells.

We then examined apical cells areas in WT and *arm>wg* embryos: these are remarkably similar between non-boundary cells and cells adjacent to either endogenous or ectopic boundaries (Fig. 5 G, H). Moreover, sampling the sectional areas throughout the 3D volume, we could not find any significant differences, suggesting that there are no significant changes in cell shapes between non-boundary and boundary cells and that apical cell areas are similar to more basal sectional cell areas (Not shown and Fig. 5G’, H’). We conclude that the boundary cells at endogenous or ectopic PSBs do not undergo apical constriction prior to epithelial folding.

Having ruled out AJs lowering and apical constriction as possible mechanisms for the role of Baz in PSBs epithelial folding, this suggests that a novel mechanism is at play. Several possibilities will be considered in future: one is that Baz could affect E-cadherin stability at PSBs (Bulgakova and Brown, 2016; Coopman and Djiane, 2016). Although E-cadherin levels are depleted at DV-oriented junctions during germ-band extension, we find in this study that at later stages, E-cadherin levels are the same at boundary and non-boundary cell-cell contacts (Fig. 1 E and Fig. 2 F). However, a possibility is that E-cadherin turnover could be different at endogenous versus ectopic boundaries because of the difference in Baz levels. This could affect in turn the mechanics of epithelial deformation: perhaps slower turnover of E-cadherin as a consequence of high level of Par-3 stabilizes adherens junctions at the PSB cell-cell contacts, which then results in greater deformation (deeper folding) for a given level of actomyosin contractility. An alternative possibility is that the increase of Baz levels at PSBs changes actomyosin contractility or cell-cell adhesion, not in the plane of the tissue, but rather in the apico-basal axis of the PSB. This could for example shorten and/or straighten the PSBs lateral cell-cell interfaces, promoting deep epithelial folding. Supporting this notion, increase in actomyosin contractility at the lateral cortex of ascidian endoderm cells is important for their invagination (Sherrard et al., 2010).

### Conclusions

Our study demonstrates that epithelial folding at the PSBs is controlled by at least two semi-independent inputs. One is actomyosin contractility along the cell-cell contacts making up the PSBs: if actomyosin contractility is increased, fold depth increases (Fig. 2). The other is the level of Baz at these cell-cell contacts: folding is suppressed when Baz is depleted at endogenous boundaries, but deepens when Baz levels are increased (Fig. 4). In the wild type, epithelial folding is minimal (shallow parasegmental grooves) because Wingless signaling depletes Baz at the boundary in parallel to maintaining high actomyosin contractility at the boundary. In *arm>wg* embryos, Hedgehog signaling counteracts this activity of Wingless posterior to the endogenous boundary, resulting in the loss of depletion of Baz, which correlates with deep epithelial folding at ectopic boundaries (Fig. 3 and Fig. S2). This is reminiscent of another case of epithelial folding: later in development, from the time when the germ-band retracts, the segmental grooves form at the *posterior* edge of the Engrailed domain (the PSBs coincide with the *anterior* edge of the Engrailed domain). In contrast to parasegmental grooves, which are shallow and require Wingless signaling, segmental grooves are deep and require Hedgehog signaling (Larsen et al., 2003). Actomyosin contractility has also been shown to be involved in segmental grove formation (Mulinari et al., 2008). A possibility is that part of Hedgehog requirement for segmental groove formation could be through a role in maintaining normal levels of Par3/Baz, counteracting Wingless-dependent depletion. Further studies are needed to test this hypothesis.

A key conclusion from our study and (Liu et al., 2016) is that specific mechanisms exist that suppresses epithelial fold formation at actomyosin-rich compartmental boundaries. We identify Par3/Baz depletion as a new factor required for this suppression, and it will be interesting in future to investigate if such depletion is required at other compartmental boundaries.

---

**Figure S1:**
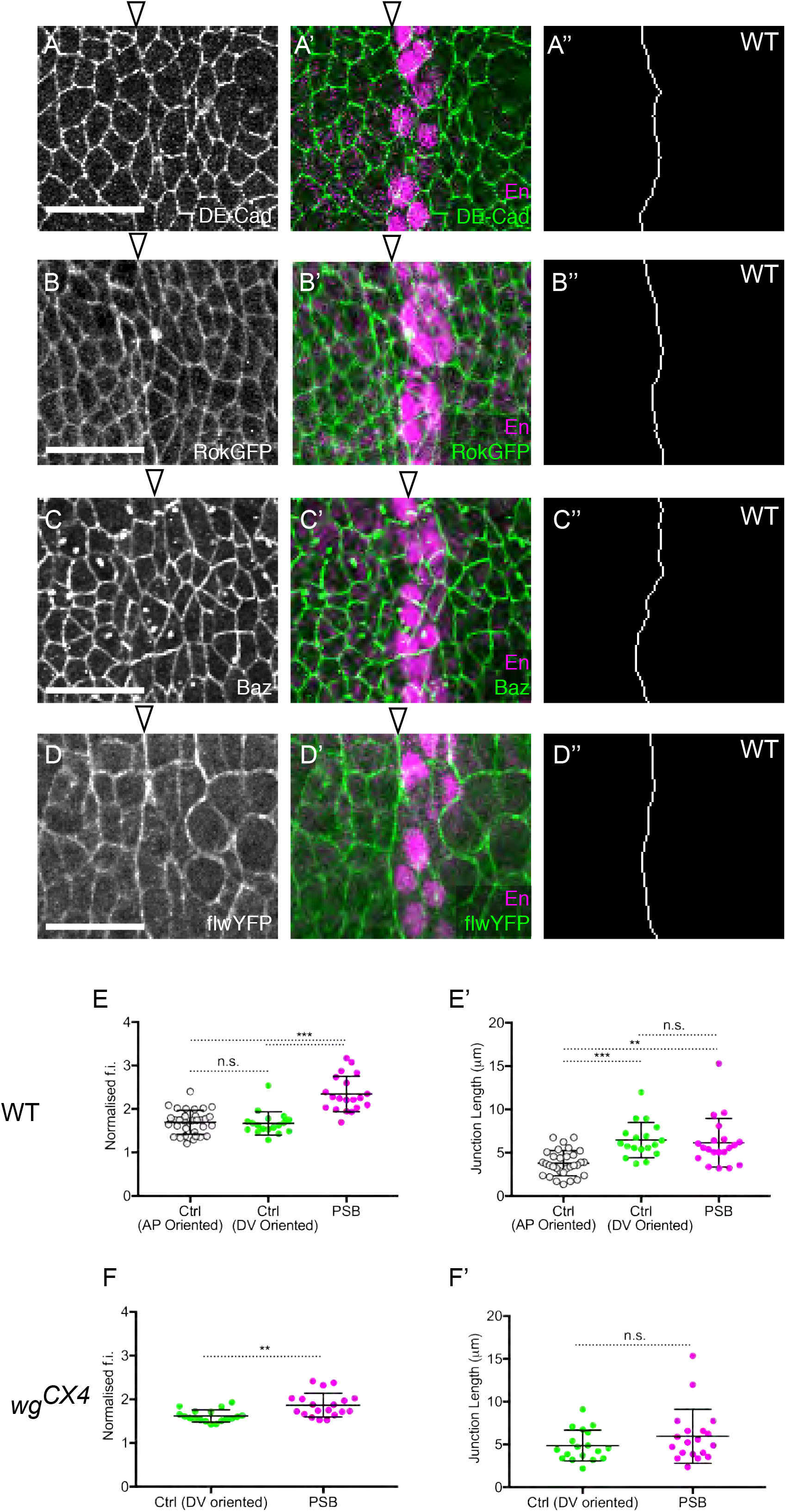
Planar polarities and actomyosin contractility at the PSBs. (A-D’’) Examples of immunostainings used for the quantification in Fig. 1E. The left panels show the immunostaining against each marker, the middle panels the merged image with Engrailed immunostaining to locate the PSBs (empty arrowheads). The right panels show the tracings of the PSB cell-cell contacts. (E-F’) Controls for the laser ablations shown in Fig. 1 F-H. (E, F) Quantification of Myosin II enrichment (using the sqh-GFP signal in live embryos) at ablated PSB and control cell-cell junctions. (E’, F’) Measurement of the length of the ablated PSB and control cell-cell junctions. DV-oriented junctions (PSB and control) are longer than AP-oriented junctions as the cells tend to be DV-elongated at this stage of development. This length difference does not appear to affect the recoil speed (see Fig. 1G).

**Figure S2:**
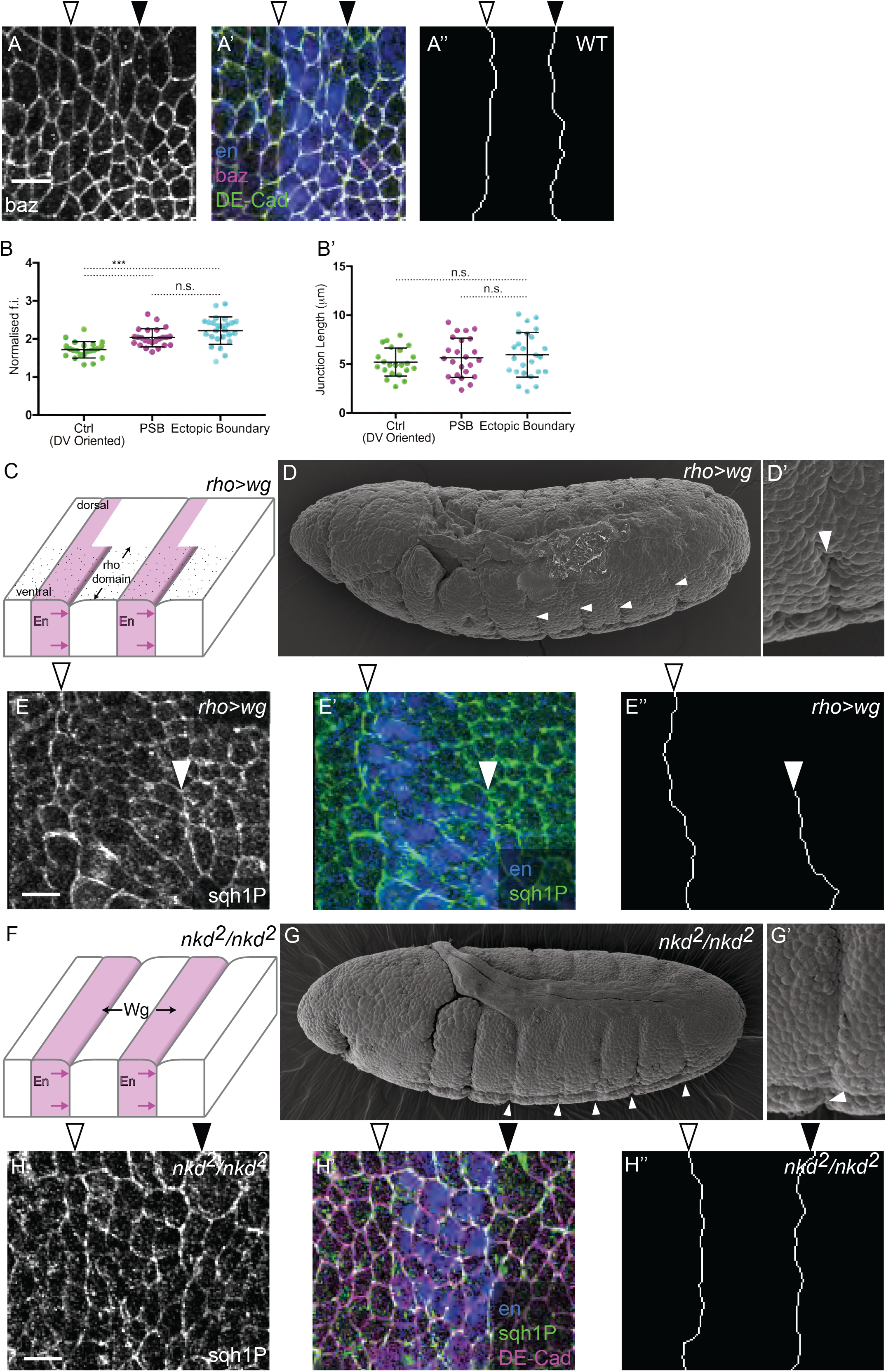
Ectopic PSBs in embryos ectopically expressing Wingless. (A-A’) Confocal images of *arm>wg* early stage 10 embryo immunostained against Baz (A), Engrailed and E-Cadherin (merge in A’). (A’’) Tracings of cell-cell contacts at endogenous and ectopic PSBs. Empty and full arrowheads label endogenous and ectopic PSBs, respectively. (B-B’) Controls for the laser ablations shown in Fig. 3D. Quantification of Myosin II enrichment (using the sqh-GFP signal in live embryos) (B) and junctional length (B’) at ablated endogenous and ectopic PSBs, and control DV-oriented cell-cell junctions. (C, E”) Formation of ectopic PSBs in embryos expressing *UAS-wg* under the control of *Rho-Gal4 (rho>wg* embryos). (C) Diagram showing the position of deep folds at ventral ectopic PSBs in *rho>wg* embryos. (D) SEM showing the short ventral folds forming at ectopic PSBs in *rho>wg* embryos (close-up in D’). (E, E’) *rho>wg* embryos immunostained against Sqh1P (E) and Engrailed (merge in E’). (E”) Tracings of the endogenous (empty arrowhead) and ectopic (full arrowhead) PSBs. (F-H”) Formation of ectopic PSBs in *nkd* null mutant embryos. (F) Diagram showing the position of deep folds at ectopic PSBs in *nkd^2^* embryos. (G) SEM showing the deep folds forming at ectopic PSBs in *nkd^2^* embryos (close-up in G’). (H, H’) *nkd^2^* embryos immunostained against Sqh1P (H), Engrailed and E-Cadherin (merge in H’). (H”) Tracings of the endogenous (empty arrowhead) and ectopic (full arrowhead) PSBs.

**Figure S3:**
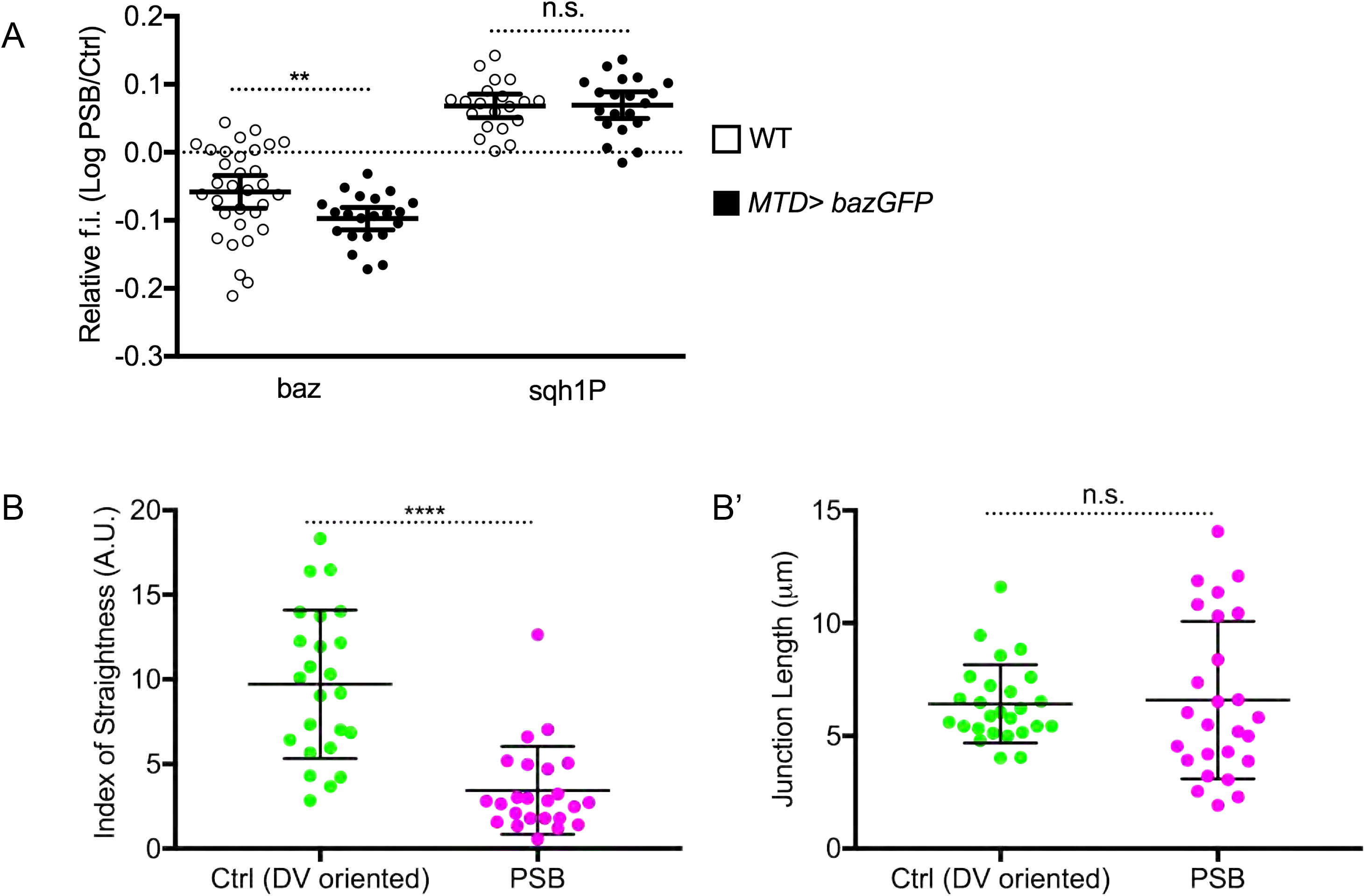
Planar polarities and actomyosin contractility at the PSBs in Baz overexpressing embryos. (A) Quantification of the enrichment of planar polarized markers at the PSBs relative to control DV-oriented interfaces in wild type (empty circles) and *MTD>bazGFP* (solid circles) stage 9 embryos. (B-B’) Controls for the laser ablations shown in Fig. 4C. Quantification of the index of straightness (B) (a proxy for junctional tension) and junctional length (B’) at ablated endogenous PSBs and control cell-cell junctions in *MTD>bazGFP* embryos.

## Material and Methods

### Fly strains

The table below lists the fly strains used and their origin. In addition, table 1 gives the genotypes used for each figure.

**Table 1:**
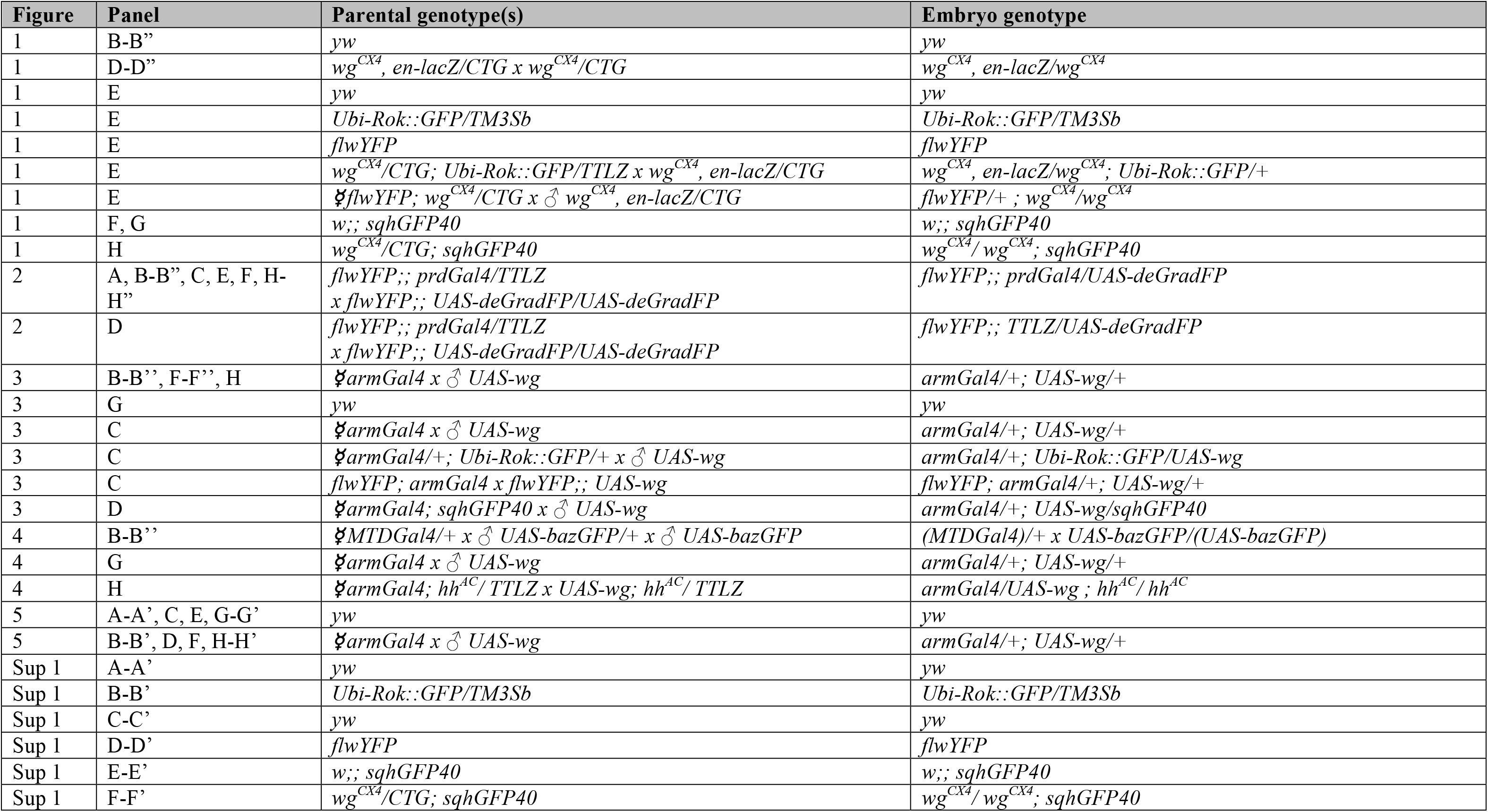
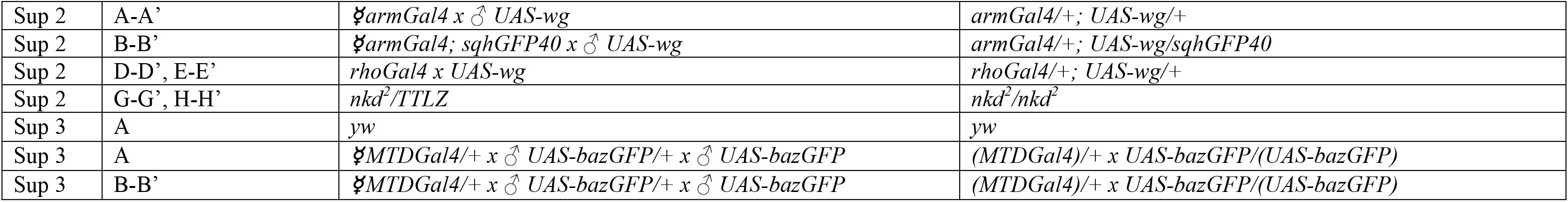
**List of genotypes used in Figures**

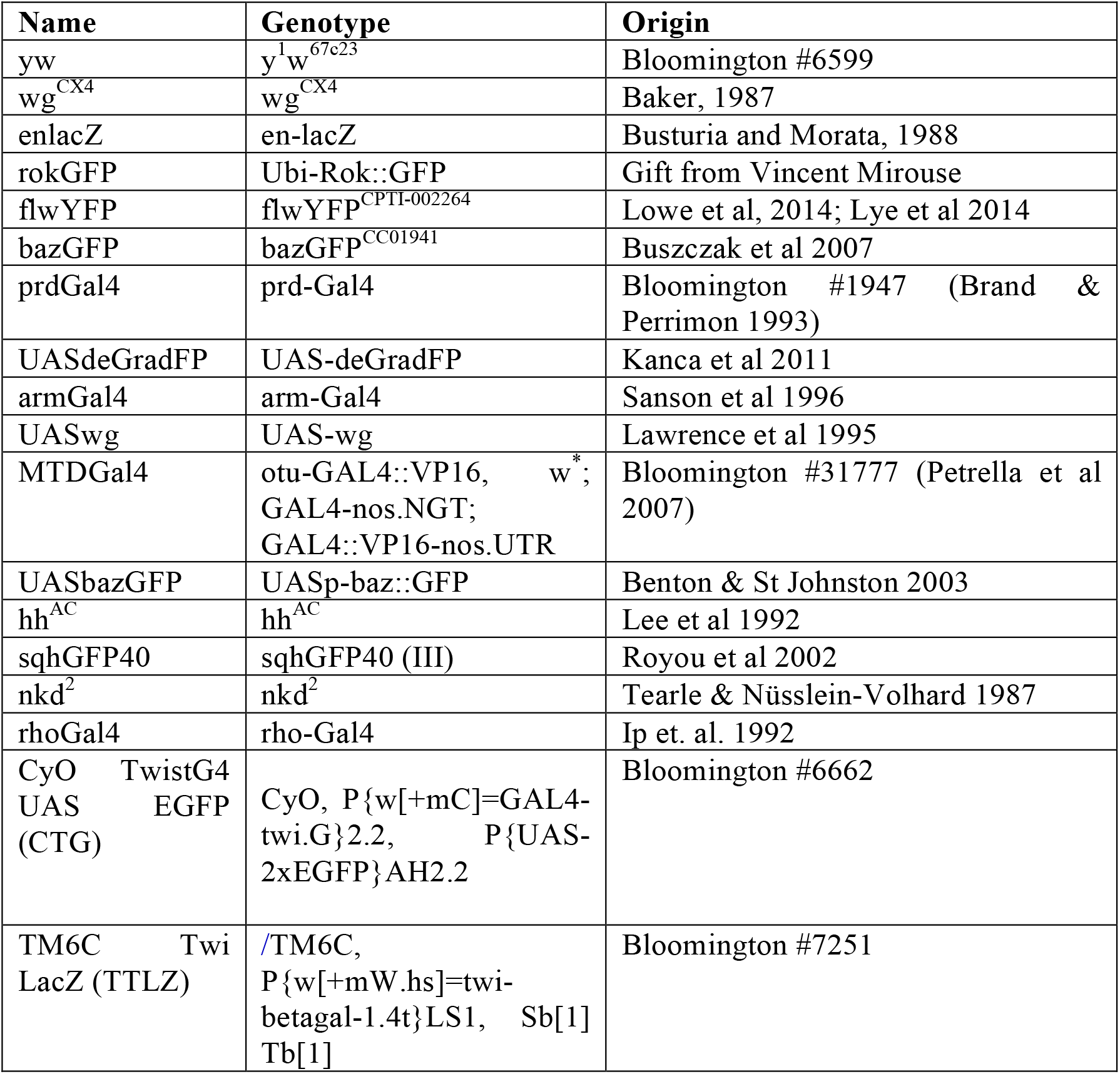

### Scanning Electron microscopy

Embryos were fixed 5 minutes in Heptane:Formaldehyde 37% (1:1) and devitellinised with Heptane:Methanol (1:1). Then, they were re-fixed immediately in 2% Glutaraldehyde, 2% Formaldehyde, 0.05M Sodium Cacodylate pH 7.4 and 2mmol/L Calcium Chloride overnight. Once rinsed twice in deionised water, embryos were treated with 1% osmium ferricyanide for 3 days. After that they were rinsed four times in deionised water, dehydrated to 100% ethanol and critical point dried. Dry embryos were mounted on carbon tabs on 12.5 mm Cambridge stubs and sputter coated with 50nm of gold. Images were taken in a FEI XL30 FEG scanning electron microscope operated at 5 kV.

### Laser ablations and analysis of recoil velocities

Laser ablation experiments were carried out on a TriM Scope II Upright 2-photon Scanning Fluorescence Microscope controlled by Inspector Pro software (LaVision Biotec) using a tuneable near-infrared (NIR) laser source delivering 120 femtosecond pulses with a repetition rate of 80 MHz (Insight DeepSee, Spectra-Physics). The laser was tuned to 927nm, with power ranging between 1.40-1.70 W. The maximum laser power allowed to reach the sample was set to 220 mW and an Electro-Optical Modulator (EOM) was used to allow microsecond switching between imaging and treatment laser powers. The laser light was focused by a 25x, 1.05 Numerical Aperture (NA) water immersion objective lens with a 2mm working distance (XLPLN25XWMP2, Olympus). Images were collected every 0.731 ms for 5 frames before the ablation and 60 frames after the ablation.

Ablations were performed during image acquisition (with a dwell time of 9.27 µsec per pixel), with the laser power switching between treatment and imaging powers as the laser was raster scanned across the sample. Targeted line ablations of ~2 µm length were performed at the centre of junctions on the PS boundary or on control, non boundary dorso-ventral (DV) oriented or antero-posterior (AP) oriented junctions, using a treatment power of 220 mW. 20-25 ablations per condition per genotype were carried out, 2-4 ablations per embryo.

To analyse recoil velocities, a kymograph spanning the ablated region was extracted using the Dynamic reslice function in Fiji, and the distance between the two ends of the cut was measured up to 30 seconds after ablation. Linear regression was performed on the first 5 timepoints after ablation and the slope of the regressed line was used a measure of the vertex recoil velocity. Statistical analysis was performed using a two tailed t-test or Mann-Whitney test.

### Immunostainings

Conventional embryonic staging was used (Campos-Ortega and Hartenstein, 1985). Embryos were collected in a basket from a one hour laying plates containing apple or grape juice hardened with agar. They were dechorionated by immersion in commercial bleach diluted 1:2 in pure water, for 2 minutes, rinsed, blotted dry and then transferred into heptane. Embryos were fixed for 5 minutes in the interface of a 1:1 solution of Heptane:Formaldehyde 37% (Fisher Scientific, Loughborough, UK) followed by manual devitelinization in PBS with 0.1% (v/v) Triton X-100 in (PTX). Embryos were then blocked PTX with 1% (w/v) bovine serum albumin (PTB) for 30 minutes, before being incubated overnight at 4ºC in primary antibody mix. They were washed three times for 15 minutes in PTX, then incubated for one hour with the appropriate secondary antibodies diluted in PTB. They were washed a further three times in PTX, and stored at -20ºC in Vectashield ® (Vector laboratories). When biotin-conjugated secondary antibodies were used an extra step was used. After the second antibody washes the embryos were incubated with streptavidin-conjugated Alexa-405 for 30 minutes before three further washes in PTX, and stored at -20ºC in Vectashield ®, Vector laboratories.

### Antibodies

Primary antibodies obtained from Developmental Studies Hybridoma Bank were:

Mouse anti-En (4D9; 1:100), Rat anti-DE-cadherin (DCAD2; 1:50), Mouse anti-Dlg (4F3; 1:500). Other primary antibodies were: Rabbit anti-Baz (1:500; provided by A. Wodarz, University of Göttingen, Göttingen, Germany), Chicken anti-ß-Gal (Abcam 1:500), Mouse anti-ß-Gal (Promega, 1:5000), Rabbit anti-β-gal (MP Biomedicals; 1:2500), Rabbit anti-Engrailed (Santa Cruz Biotechnology; 1:50), Goat anti-GFP-FITC (Abcam, 1:200), Guinea pig anti-Sqh-1P (1:100, provided by Robert E. Ward IV, University of Kansas, Kansas, USA), Mouse Phospho-Tyrosine (Cell signal; 1:100).

Secondary antibodies coupled with fluorescent dyes were obtained from Jackson ImmunoResearch Laboratories, Invitrogen and Life Technologies. Streptavidin with Alexa Fluor 405 conjugate was from ThermoFisher Scientific (1/100). Cell nuclei were stained using DAPI (4’,6-diamidino-2-phenylindole diluted in Vectashield ®, Vector laboratories, 1/500). F-actin was stained using Rhodamine-phalloidin (Molecular probes; 1:1,000).

### Confocal imaging

Embryos were mounted individually on slides in Vectashield ® (Vector laboratories) under a coverslip suspended by a one-layer thick magic tape (Scotch) bridge on either side. This flattened the embryos sufficiently so that all cells were roughly in the same z-plane. Embryos were imaged on a Nikon Eclipse TE2000 inverted microscope incorporating a C1 Plus confocal system (Nikon). Images were captured using Nikon EZ-C1 software. Optical z-stacks were acquired with a depth of 0.25 µm between successive optical z-slices. All embryos were imaged using a violet corrected 60x oil objective lens (NA of 1.4). The gain and offset were optimized for each embryo.

### Quantification of enrichment at PSBs

Two stages were used for cortical quantification: stage 10 embryos in all genotypes but *arm>wg*, which late stage 9 embryos were analyzed. Cortical signal of different proteins/markers was quantified on line traces that connect cell interfaces. The lines were manually traced by using the FIJI plugin Simple Neurite Tracer (Longair et al., 2011) and the ImageJ plugin NeuronJ (Meijering et al., 2004) based on membrane marker stainings and avoiding dividing cells. Average fluorescence intensity was quantified for 3-pixel wide line traces using ImageJ (Abramoff, 2004). Values lower than the modal pixel intensity were subtracted as background fluorescence. PSB and Ectopic Boundary interface fluorescence intensity was then normalised to En interface fluorescence intensity on a per PSB basis in all markers studied but Baz, which it was normalized to random DV tracks. Statistics were performed in Prism (GraphPad). Data from all quantifications are reported as mean ± 95% confidence intervals. Results were considered significant when P < 0.05 (*=P < 0.05, **=P < 0.02, ***=P < 0.001).

### Quantification of AJs position in the apico-basal axis and cell volume

Quantification of cell volume and adherens junctions position in the apico-basal axis. Embryos were stained with Engrailed and E-Cadherin antibodies as well as Rhodamine-Phalloidin to mark PSBs, adherens junctions and actin respectively. Then, embryos were mounted under a coverslip suspended by a two-layer thick Scotch bridge on either side. These samples were imaged on a Leica TCS SP8 confocal microscope (CAIC, University of Cambridge). Optical z-stacks were acquired with a depth of 0.33 µm between successive optical z-slices, which is the optimal z interval thickness of the 63X objective used. The gain and offset were optimized for each embryo. Fluorescence images were segmented using Real-time Accurate Cell-shape Extractor (Stegmaier et al, 2016)). Cell top was detected by the apical medial actin enrichment while cortical actin decorated cell contour. Segmented images were used in ImageJ to manually select cells of different populations (Control, PSB and ECT) in embryos of different genetic backgrounds. Selected cells were saved as region of interests and used to quantify cell area per stack and 3d render. Custom written Matlab scripts computed cell areas for the chosen cells in each plane of the stack. For the adherens junctions apico-basal position analysis contours were generated as described above in the quantification of protein enrichments at PSBs and saved as 2D binary masks. The cell walls corresponding to the regions of interest were determined by propagating these contours as open snakes on the cortical Phalloidin channel intensities (Shemesh and Ben-Shahar, 2001). These cell walls were then used to quantify the distance between the adherens junctions (E-cadherin) and the top of the cell, detected by medial actin (Phalloidin). The positions of the adherens junctions were given by the maxima of E-cadherin channel values in z direction along the wall. An estimate of the top of the cell was obtained by segmenting the Phalloidin channel stack in 2d (xz direction) via robust statistics based thresholding of the wavelet coefficients of the image. 2d projections of intensities in the E-cadherin and Phalloidin channel (across the width of the bounding box for each input contour) were saved as a mean of quality control by visual inspection. The distance between adherens junctions and the closest point of the cell top was computed taking into account voxel anisotropy. Finally, as a post-processing step of removing outliers, the highest 10% of distances were discarded for each region.

